# Sex-dependent effects of CYP46A1 overexpression on cognitive function during aging

**DOI:** 10.1101/2021.04.23.441050

**Authors:** María Latorre-Leal, Patricia Rodriguez-Rodriguez, Luca Franchini, Makrina Daniilidou, Francesca Eroli, Bengt Winblad, Kaj Blennow, Henrik Zetterberg, Miia Kivipelto, Manuela Pacciarini, Yuqin Wang, William J. Griffiths, Ingemar Björkhem, Anna Sandebring Matton, Paula Merino-Serrais, Angel Cedazo-Minguez, Silvia Maioli

**Author notes:** Corresponding author: Silvia Maioli, PhD, Karolinska Institutet, Department of Neurobiology, Care Sciences and Society. Center for Alzheimer Research, Division of Neurogeriatrics, Akademiskastråket 1, Bioclinicum, J9:20, 17146 Stockholm, Sweden.

## Abstract

Cholesterol turnover and CYP46A1 regulation are reported to be crucial for memory functions. An increasing body of evidence shows that CYP46A1 activation is able to reduce Alzheimer’s Disease (AD) pathological processes. In this study we report for the first time that CYP46A1 overexpression and increase of 24S-hydroxycholesterol (24OH) induces sex-specific changes in synaptic functions in aged mice, being beneficial in females while detrimental in males. The positive effects on cognition in aged CYP46A1 overexpressing female mice were accompanied by morphological changes in dendritic spines and enhancement of estrogen receptor signaling in hippocampus. In aged males, CYP46A1 overexpression leads to anxiety-like behavior and worsening of spatial memory, followed by decreased dendritic spine density and higher 5α-dihydrotestosterone (DHT) levels in hippocampus. Further, analysis of cerebrospinal fluid (CSF) from AD, mild cognitive impairment and healthy patients revealed that 24OH was negatively associated to markers of neurodegeneration in women but not in men. Based on our results, CYP46A1 activation may represent a pharmacological target that could specifically enhance brain estrogen receptor signaling in women at risk of developing AD. Finally, this study highlights the importance of taking into account the sex-dimension in both preclinical and clinical studies of neurodegenerative diseases like AD.

## INTRODUCTION

In the brain, the neuron-specific enzyme Cholesterol 24-Hydroxylase (CYP46A1) promotes cholesterol excretion by converting cholesterol into its oxidized metabolite, 24(S)-hydroxycholesterol (24OH) ^1^. The maintenance of a continuous flux of cholesterol into 24OH is essential for brain function, including learning and memory ^2^. CYP46A1 Knock Out (KO) mice display memory deficits ^3^, while old females overexpressing CYP46A1 showed better spatial memory performances and increased levels of synaptic markers and other related memory proteins ^4^. On the other hand, it has been reported that up-regulation of CYP46A1 may negatively affect aging processes, where stress conditions could lead to hyperactivity of CYP46A1 and cholesterol loss ^5, 6^.

Increasing body of evidence suggest CYP46A1 activation is able to reduce Alzheimer’s Disease (AD) pathological processes ^7, 8^. Adeno associated vector (AAV) injections encoding CYP46A1 in hippocampus of AD mouse models reduced amyloid-beta (Aβ) burden and restored spatial memory performances ^9, 10^, whereas CYP46A1 inhibition led to opposite effects ^11^. Pharmacological activation of CYP46A1 obtained by chronic administration of the HIV-drug Efavirenz (allosteric modulator of CYP46A1) reduced glial activation and rescued memory impairment ^12^. Recently, decrease of Aβ secretion and Tau phosphorylation was described in human iPS-derived neurons from patients with familiar AD that were treated with Efavirenz and statins^13^. Levels of the oxysterol 24OH are modified during AD progression. AD post mortem brains showed significant reduction of 24OH and CYP46A1 ^6, 14^, although in some studies cerebrospinal fluid (CSF) levels are found to be increased ^15, 16^. In addition to AD, CYP46A1 activation revealed neuroprotective effects in preclinical models of Huntington’s Disease ^13, 17^ and Parkinson’s Disease ^18^.

Oxysterols are not only passive derivates of cholesterol, but they interact with different receptors in the periphery and in the central nervous system (CNS). In particular, 24OH was previously reported to affect brain metabolism and cognitive function by interacting with Liver X Receptor (LXR), N-Methyl-D-Aspartate receptor (NMDAR)^19, 20^ and Retinoic Acid-related orphan receptor α and γ (RORα and RORγ) ^21^. Interestingly, LXRα and β activations promote the production of neuroactive steroids in the brain (neurosteroidogenesis)^22^, and oxysterols as 27 hydroxycholesterol (27OH) were described as selective modulator of estrogen receptors^23^.

A significant higher number of women are suffering from AD^24^ and it has been shown that women convert faster from mild cognitive impairment (MCI) to AD compared to men ^25^. Nevertheless, the reasons behind sex-differences in AD pathogenesis and progression have been largely unexplored ^26^. Considering the relevance of CYP46A1 as a possible target to cure AD and that cholesterol metabolism could differently affect men and women, we aim to investigate the effects of CYP46A1 activation in cognitive functions and neurodegeneration in a sex-specific manner.

Here, we describe a novel role of 24OH in brain as modulator of 17β-estradiol (E2) and neurosteroid signaling in neurons through nuclear receptors pathways. Estrogen signaling was found enhanced in hippocampus from old female mice overexpressing CYP46A1. Differently from females, old CYP46A1 overexpressing male mice showed increased levels 5α-dihydrotestosterone (DHT) levels in hippocampus. We report that overexpression of CYP46A1 in vivo induces sex-specific changes in synaptic functions in aged mice, being beneficial in females while detrimental in males. Finally, analysis of cerebrospinal fluid (CSF) from memory clinic patients show that 24OH levels were negatively associated to markers of neurodegeneration in aged women but not in men.

## MATERIAL AND METHODS

### Transgenic mice

Female and male CYP46A1 HA-tagged heterozygous transgenic mice (Cyp46 Tg) and wild type littermates (Tg-) were studied at 9, 16 and 20 months of age. All mice were kept under controlled temperature (21± 1°C) and humidity (55 ± 5%) on a 12-h light-dark cycle and food/water were provided ad libitum. Experimental procedures were conducted in accordance with the European regulation and approved by the local Ethical Committees at Karolinska Institutet, Stockholm. All the efforts were made to minimize the suffering of the mice.

### Rat primary culture and treatments

Hippocampal tissue was isolated from E18 Sprague-Dawley rat embryos (Charles River). Neurons in primary culture were seeded (125 000 cells/cm^2^) in Neurobasal® media (Thermo Fisher), supplemented with 2% B-27® (Thermo Fisher), 2 mM GlutaMAX^™^ (Thermo Fisher), 100 units/ml penicillin and 100 μg/ml streptomycin (PEST, Thermo Fisher), and incubated at 37°C in a humidified 5% CO_2_-containing atmosphere. All experiments were performed after 10 days in culture. For treatments, media was removed and replaced with Neurobasal® media with 1 μM of 100% 24S-hydroxycholesterol (Instruchemie, Netherlands) prepared in ethanol and 10 nM DHT (Sigma-Aldrich) resuspended in methanol. Media with the same final concentration of vehicles EtOH/MeOH was used as a control. The cells were then incubated for different time points at 37°C in a humidified 5% CO_2_-containing atmosphere.

### Behavioral tests

At 20 months of age, the mice were tested for the following behavioral tasks: Elevated Plus Maze (EPM), Dark/Light Box (DLB) and Y-maze test, while mice at 9 and 16 months of age were tested for Morris Water maze (MWM). The order of the different tests was chosen based on the level of stress associated with the procedure^28^. Mice were placed in the experimental room 30 minutes for habituation to the new environment and all the tests were run between 9:00 and 15:00. The experiments were performed in white light and by the same experimenters. EPM, DLB and Y-Maze protocols were run as previously described in details ^29^. For MWM test, the mice were trained to find a hidden escape (the platform) as previously described ^4,16^. The different parameters from these tests were assessed with the video-tracking system Ethovision XT 14 (Noldus, Netherlands), connected to a video camera placed above the equipment.

### Golgi staining and dendritic spine analysis

After behavioral testing, five 20 months old Cyp46 Tg and Tg-female and male mice were randomly selected and processed for Golgi analysis. Left brain hemispheres were stained with the Golgi–Cox solution using the FD Rapid GolgiStain^™^ Kit (NeuroTechnologies Inc., Columbia, MD, USA) according to manufacturer’s instructions. The impregnated hemispheres were then rapidly frozen and embedded in Tissue Freezing Medium before sectioning on the cryostat (at −20 °C/-22 °C). Coronal sections of 150□μm of thickness were mounted on gelatin-coated microscope slides with Permount Mounting Medium (Thermo Fisher Scientific) and dried at room temperature overnight. The following day silver staining was performed.

A light microscope (Nikon Eclipse E800) was used to acquire the images with 100x objective, oil immersion (NA: 1.30). Eight to ten slices per brain containing the whole hippocampus were selected for the analysis. Images were collected in the stratum radiatum from the CA1 hippocampal region. Only collateral dendrites (stratum radiatum) from the pyramidal neurons were chosen and dendrites from 2-3 orders, were analyzed. The cells to analyze were selected based on the following criteria: a) neurons were relatively complete (3 order or greater of dendrites were visible); b) they were fully impregnated with staining; and 3) there was minimal or no overlap with other labeled neurons. A total dendritic length of 300 μm per animal was analyzed. All analyses were conducted blinded to genotype. The analysis of the images was performed with ImageJ software (NIH, US).

### Immunoblotting

Hippocampal mouse tissue and hippocampal rat neurons in primary cultures were lysed in RIPA buffer (50 mM Tris, 150 mM NaCl, 1% Triton X-100, 0.1% SDS) supplemented with protease and phosphatase inhibitor cocktail (Sigma, Aldrich 1:100, 1 mM EDTA and 1 mM EGTA). Western blot was performed as previously described ^29^. The following primary antibodies were used: anti-RAR alpha (Abcam ab28767, 1:500), anti-ER alpha (Abcam ab3575, 1:1000), anti- ER-beta (Santa Cruz Biotechnology sc-53494, 1:1000), Arc (Santa Cruz Biotechnology sc-15325, 1:1000), Aromatase (Abcam 18995, 1:1000), phospho-CREB (Cell Signaling 86B10, 1:1000), ADAM 10 (Abcam ab1997, 1:500). β-actin was used as protein loading control (Sigma A2066,1:10000). Secondary antibodies anti-rabbit, anti-mouse or anti-goat IgG were used at a 1:10000 dilution (LI-COR Biosciences GmbH, Germany). Immunoreactivity was detected by infrared fluorescence with LI-COR® Odyssey® system (LI-COR Biosciences, USA) and quantified with ImageJ software (NIH, US) by densitometry analysis of the immunoreactive bands.

### Nuclear fractionation

We used NE-PER kit (Pierce, Rockford, IL, USA) to isolate the nuclear and cytosolic fractions from hippocampal cultured neurons. The procedure was performed following the manufacturer’s protocol. Protease inhibitor cocktail (1:500, Sigma-Aldrich) was added freshly. To evaluate the purity of the fractions, immunoblotting with antibody against the nuclear marker Lamin-A/C was performed (Sigma-Aldrich SAB4200236,1:1000).

### RNA Extraction and real-time PCR (RT-qPCR)

Isolation of total RNA from rat hippocampal primary neurons was carried out as previously described ^30^. For brain tissue, mouse hippocampi were first homogenized with Trizol (Thermo Scientific), followed by the protocol mentioned above. Relative mRNA copy numbers of genes were measured by real-time PCR using Taqman Universal MasterMix (Applied Biosystems) and the following primers: *Cyp19a1, Ers1, Ers2, Arc, Bdnf, Apod, Sdr5a1, Hsd17b10, Hsd17b1, Rara, Cyp26b1, Rxrg* (Life Technologies, CA, USA). ΔCt levels for *Gapdh* were selected as control to normalize ΔCt levels for each sample.

### Enzyme-linked immunosorbent assay (ELISA)

ELISA for quantitative analysis of DHT and E2 (ELISA Kit LBio and Abcam respectively) in mouse brain tissue was adapted to the manufacturer’s instruction. 20 mg of dissected hippocampus was homogenized in lysis buffer (Tris-HCl 50 mM, NaCl 150 mM, 1% Triton, adjusted to pH 7.4). Methanol was added for extraction of steroidal hormones. After incubation and centrifugation, the supernatant was transferred to a new tube, dried and resuspended in the ELISA-compatible buffer solution.

### Memory clinic patient population

The study included 90 patients diagnosed with either subjective cognitive decline, mild cognitive impairment or Alzheimer’s disease including a total of 43 men and 48 women. This population includes only subjects that were not using medications for diabetes, hypercholesterolemia and hypertension. The patients were examined at the Karolinska University Hospital memory clinic in Huddinge, Sweden. The study was approved by the Regional Ethical Review Board in Stockholm, and written informed consent was obtained from all patients.

### CSF biomarkers

CSF samples were collected by standard lumbar puncture between the L3/L4 or L4/L5 intervertebral space using a 25-gauge needle. CSF was aliquoted in polypropylene tubes and stored at −80°C until further analysis. CSF Aβ42, t-tau, and p-tau concentrations were measured on fresh samples with commercially available sandwich enzyme-linked immunosorbent assays (Innogenetics, Ghent, Belgium) according to standardized protocols in the clinic. 24OH was analyzed as the sum of the esterified and free molecule by liquid chromatography and mass spectrometry as previously described ^31^ but with the addition of a base hydrolysis step in 0.35 M KOH prior to solid phase separation of total oxysterols from cholesterol. Neurofilament light chain (NFL) was measured as reported previously ^32^.

### Statistical analysis

Data are expressed as mean ± standard error of the mean (SEM), with N indicating the number of mice or experiments performed. When comparing two groups, t-Student or Mann-Whitney tests were used for parametric and non-parametric data respectively. One-way ANOVA and Two-way ANOVA repeated measurements followed by Tukey’s multiple comparison test was used to analyze data when two independent variables were present. Repeated measure way ANOVA followed by Tukey’s multiple comparison test was used for analyses of MWM test. Memory clinic population data were compared across men and women in the cohort with Mann-Whitney U tests for continuous variables and χ^2^ for categorical. Correlations were performed by Spearman analysis. Aβ42, p-tau, NFL and 24OH were log-transformed and t-tau was square root transformed to increase normality. To assess the association of 24OH with the AD and neurodegeneration biomarkers, we performed separate multiple linear regression models in pair-wise combinations between 24OH and each of the biomarker (Aβ42, t-tau, p-tau, NFL) in all participants and in men and women separately, adjusting for age and diagnosis. 24OH was set as the independent variable in all models. GraphPad Prism 7 software (San Diego, CA, USA) and IBM SPSS 27 (SPSS Inc., Chicago, IL) were used and a P value <0.05 was considered as significant.

## RESULTS

### CYP46A1 overexpression induces sex-specific changes in behavior and CA1 dendritic spine morphology in aged mice

A battery of behavioral paradigms was used to evaluate anxiety-like behavior, hippocampal-dependent learning and memory of old male and female Cyp46 Tg mice and their age and sex-matched wild type littermates Tg-, as shown in Figure 1A-H. During the EPM test, 20 months old Cyp46 Tg female mice displayed no differences in the percentage of time spent in open arms when compared to the control group (Figure 1A). When animals were assessed for the Y-maze task, Cyp46 Tg alternated at significantly higher levels than the chance level of 50% (P=0.022, One sample T-test, Figure 1B), while the control group did not so, indicating an enhancement of spatial working memory in Cyp46 Tg mice. Moreover, a significant difference was found when mice were tested for the dark-light box test, with Cyp46 Tg mice displaying a significant lower latency time to enter the lit compartment (P=0.0149, Mann Whitney test) in comparison to Tg-mice (Figure 1C), suggesting a decrease of anxiety-like behavior in female mice overexpressing CYP46A1. 20 months old male mice were also tested for EPM and Y-maze test. Oppositely to Cyp46 Tg females, Cyp46 Tg male mice spent a significantly shorter time in the open arms of the EPM when compared to their age-matched littermates (P=0.0047, Mann Whitney test, Figure 1D), indicating that overexpression of CYP46A1 in old male mice lead to increase of anxiety-like behavior. During the Y-maze test, no significant differences were found between groups however it was possible to observe a higher number of Cyp46 Tg mice that alternated under the chance level of 50% as compared to Tg-mice (Figure 1E), suggesting a possible impairment of spatial working memory induced by CYP46A1 overexpression in males. Hippocampal-dependent spatial learning and memory was tested in 16-month-old mice by using Morris water maze task, as illustrated in Fig. 1F, 1G and 1H. In the learning phase from day 1 to day 4, no differences between groups were found in the latency time to reach the hidden platform (Two-way ANOVA, Repeated measurements, effect of days: P=0.0094, F= 4,634, Figure 1F), indicating that spatial learning was not affected and both groups of mice were able to locate the platform among the days. During the probe test on day 5, Cyp46 Tg male mice spent significantly shorter time in the sector where the platform was located as compared to Tg-mice (P=0.0152, Mann Whitney test, Figure 1G), indicating lower retention of the spatial information previously acquired. On the opposite, probe test heat maps showed that Cyp46 Tg female mice spent more time in the platform sector in comparison to control mice as previously reported ^4^ (Fig H). Importantly, when 9-month-old male and female mice were tested for EPM and MWM tests no differences were found between groups (Figure S1A-F).

**Figure 1.**
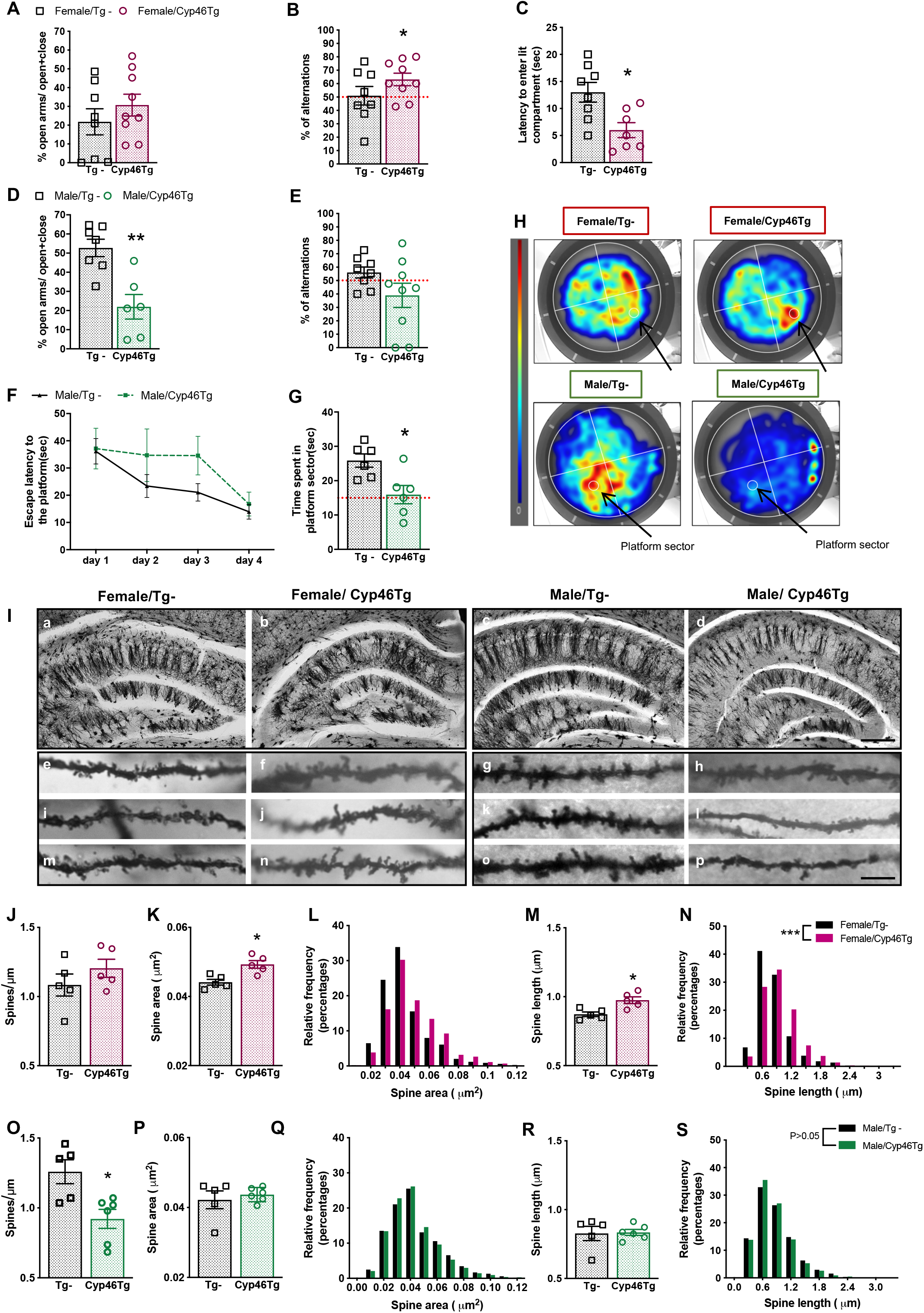
CYP46A1 overexpression enhances spatial memory and hippocampal dendritic spine area and length in aged female mice, while it leads to negative effects in males. Battery of behavioral tests in 20 months old Cyp46 Tg and Tg-female (burgundy graphs, **AC**) and male mice (green graphs, **D-G**), N=6-10 mice/group/sex. Percentage of time spent in open arms over the 5 min duration of the EPM test **(A, D,** **P<0.01). Percentage of spontaneous alternations during Y-maze test **(B, E,** *P<0.05). First latency to enter the light compartment in DLB test **(C,** *P<0.05). Escape latency (time required to reach hidden platform) over 4 days acquisition phase in MWM test (**F**), time spent in the quadrant where the platform was located during acquisition phase between both groups during probe testing (**G**, *P<0.05). Heatmaps from MWM probe test **(H)**: the occupancy rate is graded by a color map ranging from cold to warm colors (in red color the most visited zones). Representative Golgi staining images **(I)** from hippocampus **(a-d**, 4X magnification) and dendrites in stratum radiatum from CA1 region of hippocampus **(e-p**, 100X magnification) of Cyp46 Tg female mice (right panel) and male mice (left panel) and their corresponding age-matched controls Tg- (N=5-6 mice/group/sex); scale bar 1:500 μm. Analysis of dendritic spines from CA1 dendrites **(J-S)**: data are shown as dendritic spine density measured as spines/μm (**J, O,** *P<0.05), spine area in μm^2^ (**K, P**, *P<0.05), spine length in μm (**M, R**, *P<0.05), frequency distribution of spine area (**L, Q**, *P<0.05) and frequency distribution of spine length (**N, S**, ***P<0.001) from apical dendrites of Cyp46 Tg and Tg-male and female mice. All data are represented as mean ± SEM.

To elucidate whether the behavioral changes observed in mice were accompanied by alteration in dendritic spine morphology, we examined possible morphological changes in dendritic spines from apical dendrites in Golgi-stained pyramidal neurons of 20 months old Cyp46 Tg mice. In female mice, while spine density (measured as number of spines/μm) was similar between transgenic and wild type groups (Fig 1J), dendritic spines of apical neurons in CA1 region were significantly bigger and with longer area in Cyp46 Tg females as compared to their littermates (Figure 1K, 1M, Mann Whitney test P=0.0159 for both area and length). Moreover, Cyp46 Tg female spine distribution displayed higher frequency in bigger size and length in comparison to Tg-(two-sample Kolmogorov-Smirnov test, P<0,0001, Figure 1L, 1M). Cyp46 Tg males presented a significant lower spine density in CA1 pyramidal neurons when compared to their controls (Mann Whitney test, P=0.0108, Figure 1O). Frequency distribution of area and length of Cyp46Tg males showed a reduction on size and length of dendritic spines as compared to the control group (two-sample Kolmogorov-Smirnov test, P=0.0480 and P=0.0162 respectively, Figure 1Q, 1S).

### Sex-specific changes in estrogen receptors levels and neurosteroid metabolism in the hippocampus of Cyp46Tg aged mice

Considering the sex differences observed in old Cyp46Tg mice, we next aimed to clarify whether these divergent effects might be mediated by neuroactive steroids. The sexual hormones E2 and DHT, can be locally synthetized in the brain, using cholesterol as the main substrate (Figure 2O) ^33^. We first analyzed hippocampal levels of E2 and DHT in males and females Cyp46 Tg by ELISA. E2 did not change between groups in females (Figure 2M). However, DHT levels were significantly increased in hippocampus of males Cyp46Tg as compared to their control littermates (Figure 2N, Mann Whitney test, P=0.0451).

**Figure 2.**
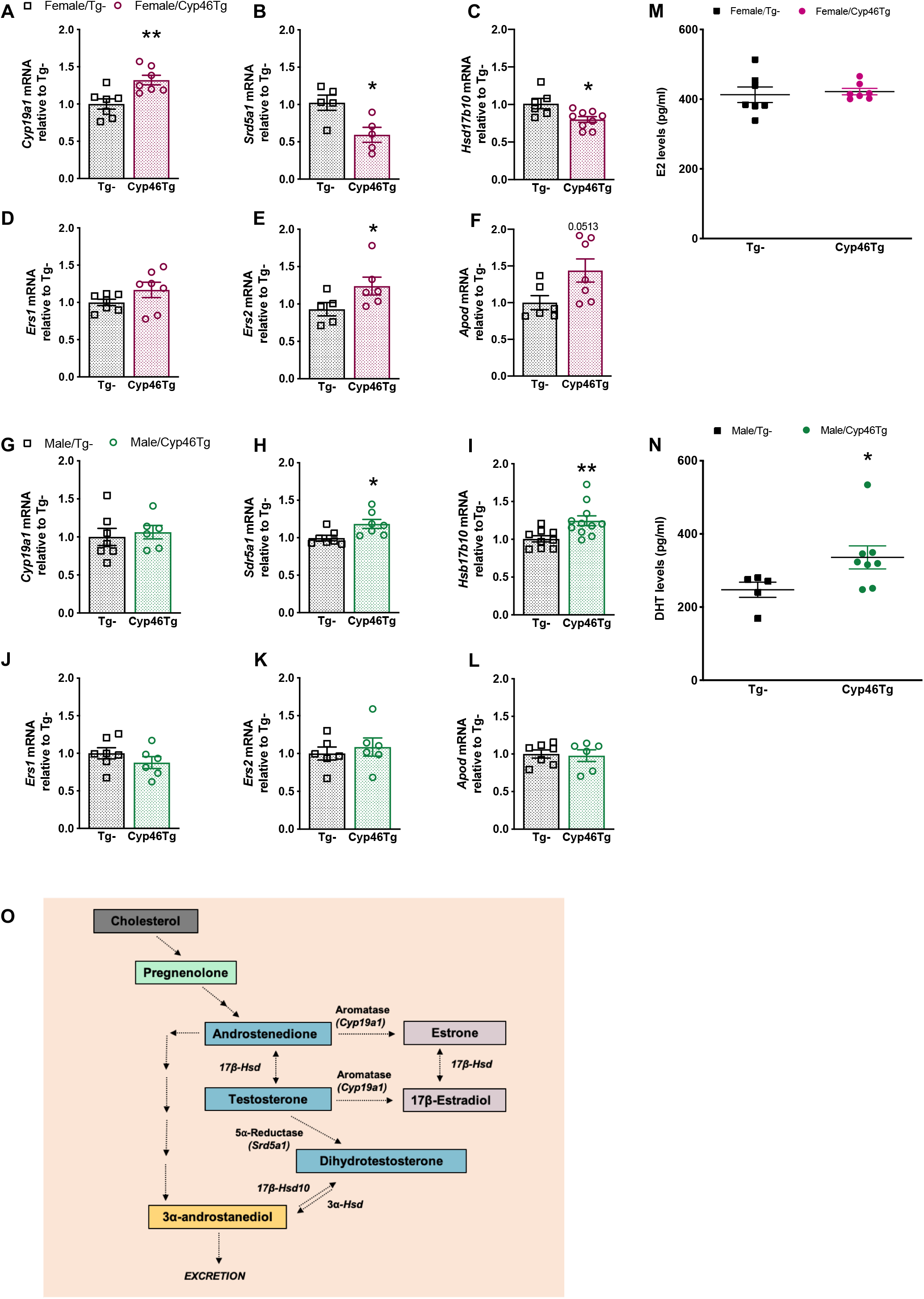
Estrogen receptor signaling is increased in hippocampus of old Cyp46 Tg female mice, while DHT levels and DHT-related signalling are higher in transgenic males. mRNA expression levels of *Cyp19a1, Srd5a1, Hsd17b10, Ers1, Ers2 and Apod* genes measured by RT-qPCR in hippocampus from 20 months old Cyp46 Tg female (**A-F**) and male mice (**G-L**) in comparison to their Tg- controls. All mRNA levels are normalized by *Gapdh* mRNA levels (*P<0.05, **P<0.01, N=5-11 mice/ group/sex). ELISA measurements of E2 levels in hippocampus from Cyp46 Tg and Tg- female mice **(M)** and DHT levels in hippocampus from Cyp46 Tg male mice **(N)** compared to controls (*P<0.05, **P<0.01, N=5-8 mice/group/sex). All data are presented as mean ± SEM. **(O)** Representative scheme for the neurosteroids synthesis and metabolism starting from cholesterol as a precursor molecule in brain.

Neuroactive steroids synthesis pathway in the brain is tightly regulated by different enzymes as shown in the schematic figure 2O. We measured mRNA expression levels of key-enzymes and receptors involved in E2 and DHT biosynthesis. The expression levels of aromatase (*Cyp19a1*) were significantly increased in the hippocampus of old Cyp46Tg female mice as compared to Tg-mice (Mann Whitney test, P=0.0070, Figure 2A). Nevertheless, *Cyp19a1* mRNA levels remained unchanged in old male Cyp46Tg when compared to Tg-controls (Fig 2G). Expression of 5α-Reductase (*Srd5a1*, responsible for DHT synthesis from testosterone) and Hydroxy-steroidodehydrogenase-17β-10 (*Hsd17b10*, that synthetizes DHT from 3α-Diol) was significantly decreased in transgenic female mice compared to control (Mann Whitney test, P=0.0360 and P=0.0317 respectively, Fig 2B, 2C). Contrary to the effect observed in females, levels of *Srd5a1* and *Hsd17b10* were found to be increased in Cyp46Tg males corresponding with the increase in DHT levels observed in these mice (Mann Whitney test, P=0.0175 and P=0.0042 respectively, Figure 2H, 2I). Expression levels of estrogen receptor β (*Ers2*) were significantly increased in old Cyp46Tg female mice in comparison to Tg- animals (Mann Whitney test, P=0.0411, Fig 2E). CYP46A1 overexpression did not affect *Ers1* (the gene that codifies for ERα) in old Cyp46 Tg females. Levels of Apolipoprotein D (*Apod*), target gene of *Ers2*, were also raised in Cyp46Tg females as compared to controls, with a tendency towards significance (Mann Whitney test, P=0.051, Fig 2F). *Ers1, Ers2* expression remained unchanged in old male Cyp46 when compared to Tg- animals (Figure 2J and 2L).

### 24OH activates estrogen receptors and neurosteroid metabolism through nuclear receptor signaling in hippocampal neurons

To further clarify whether estrogen signaling and neurosteroidogenesis could be affected by 24OH, we treated hippocampal neurons in primary culture with 1 μM 24OH (Figure 3). In agreement with the results observed *in vivo*, RT-qPCR analysis revealed a significant increase in mRNA levels of *Ers2* (Figure 3B, Mann Whitney test, P=0.0102) after 24 hours 24OH treatment, while *Ers1* levels were unchanged between control and 24OH-treated neurons (Figure 3A). Moreover, the enzymes coded by *Cyp19a1* and *Hsd17b1* were significantly increased in 24OH treated cells when compared to control cells (Mann Whitney test, P=0.0022, P<0.001; Figure 3C, 3D). We next analyzed ERβ and ERα protein levels, 5 hours treatment with 24OH triggered a significant raise of ERβ in the nuclear fraction of 24OH-treated neurons compared to vehicle-treated controls (Mann Whitney test P=0.0312, Figure 3K). ERα nuclear levels remained unchanged (Figure 3J). In addition, a trend towards a significance increase in treated cells was found for Aromatase protein levels compared to control cells (Mann Whitney test, P=0. 0842, Fig 3O).

**Figure 3.**
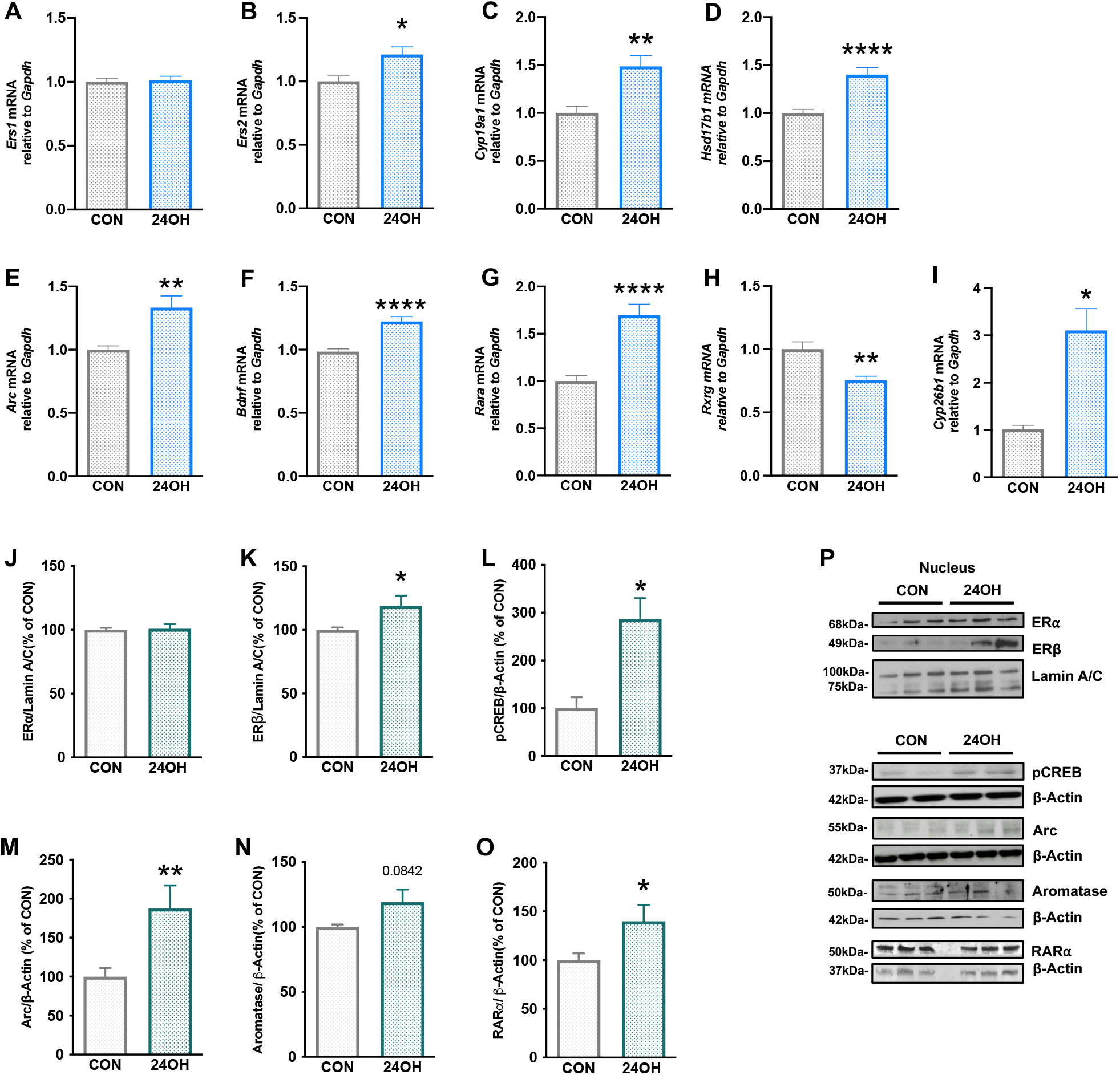
24OH in vitro treatments trigger estrogen and neuroactive steroid signaling through nuclear receptor pathways. Comparative analysis of mRNA levels of *Ers1, Ers2, Cyp19a1, Hsd17b1, Arc, Bdnf, Rara, Rxrg* and *Cyp26b1* genes, assessed by RT-qPCR in primary cultured neurons treated with 1 μM of 24OH or its vehicle (CON), as reference (**A-I**, **P<0.01, **** P<0.0001, N=3-5 independent experiments in triplicate and quantified using *Gapdh* as housekeeping gene). Western blot analysis **(J-O)** and representative blots **(P)** of ERα and ERβ, Arc, pCREB, RARα and Aromatase protein levels after 24-hour treatment with 1μM of 24OH in primary cultured neurons (*P<0.05, ** P<0.01, N=3 independent experiments, each in triplicate, were performed). Data are shown as mean ± SEM of immunoreactivity (OD × area of the band) and normalized by actin levels.

Afterwards, we assessed estrogen downstream targets, including known regulators of synaptic plasticity, such as Arc, Bdnf and phosphorylated CREB (p-CREB). RT-qPCR analysis confirmed an increase in mRNA levels of *Arc* and *Bdnf* in 24OH-treated neurons as compared to vehicle-treated cells (Mann Whitney test, P=0.0089, P<0.0001, respectively, Figure 3E, 3F). Western blot of hippocampal neurons treated for 24 hours revealed a significant increment of p-CREB (P=0.05) and Arc (P=0.070) protein levels compared to control (Mann Whitney test, Figure 3L, 3M).

To discern the mechanisms by which 24OH is able to induce estrogen signaling, we assessed the possible interplay of 24OH on nuclear receptor signaling, as LXR and Retinoic acid receptor signaling, previously known to activate neurosteroidogenesis. 6 and 12 hours treatments of neurons with 1 μM 24OH significantly increased *Srebf1*, a gene target of LXR signaling (Mann Whitney test, P=0.0400, P=0,0079, Supplementary Figure 2A). The mRNA levels of Retinoid Acid Receptor α (*Rara*), as well as its target genes *Cyp26b1* were significantly elevated in hippocampal neurons treated with 24OH for 12 hours (Mann Whitney test, P<0.0001 and P=0.0043 respectively, Figure 3G and 3I). Levels of Retinoid X Receptor γ (*Rxrg*), known to dimerize with RARα, were significantly decreased in 24OH treated cells as compared to vehicle cells (Mann Whitney test, P=0.0159, Fig 3H). In addition, a significant increment on RARα protein levels were shown when treated neurons were treated with 1 μM 24OH (Mann-Whitney test P=0.042, Fig 3O).

### DHT treatment in hippocampal neurons counteracts the effects of 24OH on estrogen and retinoic acid receptor signaling

As shown in Figure 3, 24OH activates estrogen receptor signaling and neuroactive steroid metabolism possibly by activation of nuclear receptor pathways. To assess the interplay between 24OH-induced estrogen signaling and DHT (found to be elevated in brain of Cyp46 Tg old males), we performed treatments in hippocampal neurons from rat primary culture with 10 nM DHT alone or in combination with 1 μM 24OH and (DHT+24OH). 6-hour treatment with DHT and DHT+ 24OH showed no significant changes in *Ers1 and Ers2* mRNA levels measured by RT-qPCR (Figure 4A and 4B). Levels of *Cyp19a1* were increased after the neurons were treated with DHT alone (P=0.0056), while they remained unchanged when DHT was combined to 24OH, as compared to control cells (One-way ANOVA, followed by Tukey post hoc test, Figure 4C). No significant differences were found in *Hsd17b1* levels with DHT nor DHT+24OH treatments (Fig 4D). DHT treatments did not affect *Arc* and *Bdnf* mRNA levels when compared to control cells, while treating neurons with DHT+24OH significantly increased levels *of Arc* and *Bdnf* when compared to vehicle cells (One-way ANOVA, followed by Tukey post hoc test, P=0.0006 and P=0.0167 respectively, Figure 4E and 4F). *Rara* mRNA levels, together with its target genes *Cyp26b1* and *Rxrg* were also measured in hippocampal neurons treated with either DHT or the combination of DHT+24OH. Differently from 24OH treatments (shown in Figure 3), treatments with DHT and DHT + 24OH did not augment *Rara* and *Cyp26b1* expression levels compared to vehicle cells and no differences were found between treatments (Figure 4G and 4H). DHT treatment alone or in combination with 24OH did not show significant changes in the expression of *Rxrg* as compared to controls (Figure 4I). Since RAR signaling is known to promote the expression of A Disintegrin and Metalloproteinase 10 (ADAM10) ^34^, we further assessed ADAM10 protein levels in hippocampal homogenates from old Cyp46Tg and mice and compared to controls. Cyp46Tg female mice showed enhanced hippocampal ADAM10 protein levels when compared to controls, while no difference could be inferred between groups in male mice (Mann Whitney test P=0.0286, Supplementary Figure 1G and 1H).

**Figure 4.**
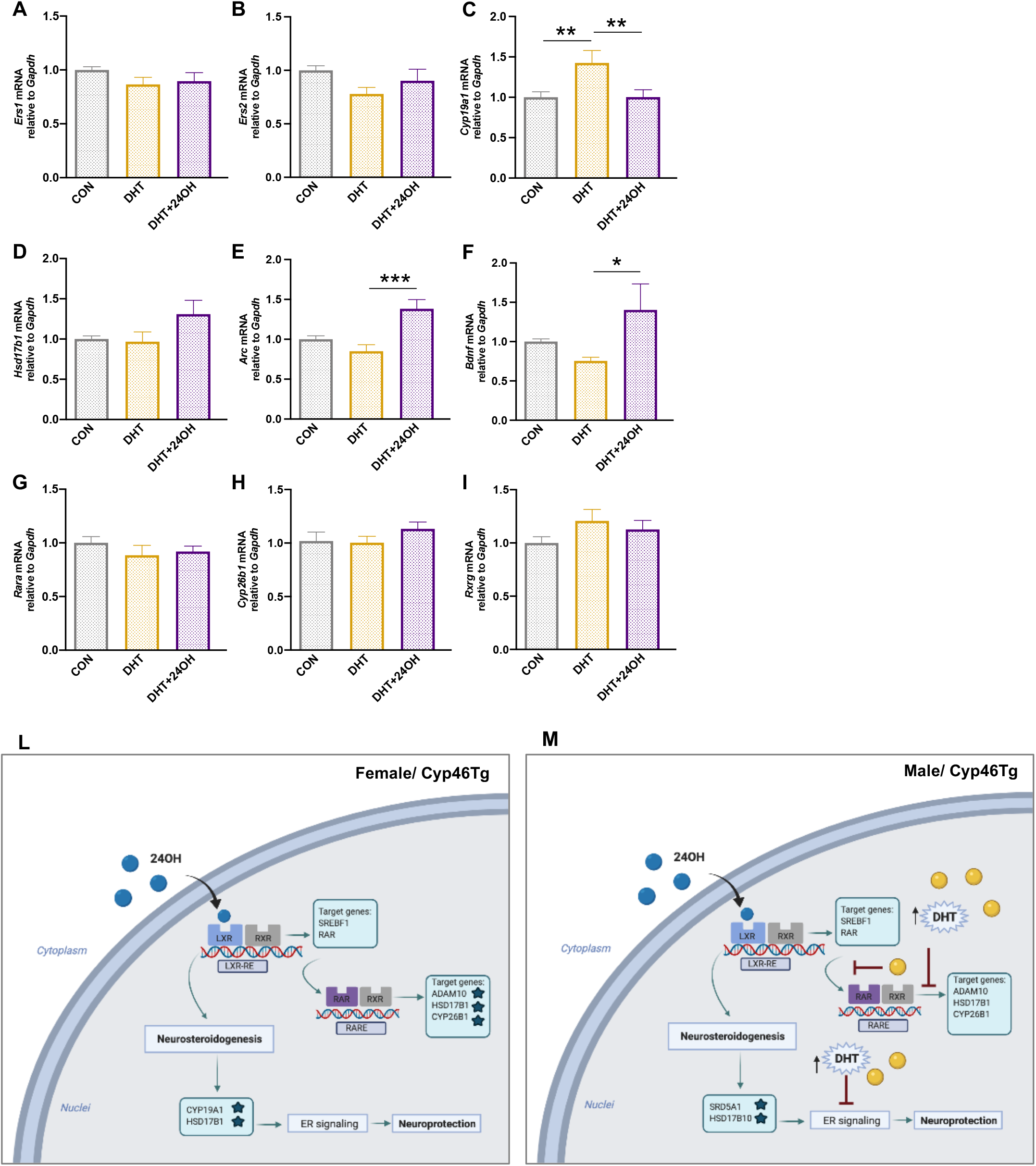
DHT treatment in neurons reverts 24OH-mediated estrogen and retinoic acid receptor activation. Hippocampal neurons were treated with 10 nM DHT and with a combination of 10 nM DHT + 1 μM of 24OH and its vehicle (CON) as reference. mRNA levels of *Ers1, Ers2, Cyp19a1, Hsd17b1, Arc, Bdnf, Rara, Rxrg* and *Cyp26b1* were measured by RT-qPCR and normalized using *Gapdh*, data shown as mean± SEM (**A -I**, *P<0.05, **P<0.01,***P<0.001, N=3 independent experiments, each in triplicate, were performed). Proposed mechanism for 24OH in Cyp46 Tg female and male aged mice **(L, M)**. In the brain, 24OH binds to LXR, activating RARE promoter and RARα signaling in neurons. 24OH induces neurosteroidogenesis via LXR, enhancing estrogen signalling to promote neuroprotection in transgenic female mice. In contrast, the presence of high levels of DHT in male mice seem to counteract 24OH effects on RARα and estrogen receptor signalling. The star symbol means significant increase of genes/proteins. Images were created with Biorender.com.

### CSF 24OH is negatively associated with tau and NFL in women but not in men, in a memory clinic cohort

We next investigated the possible relationship between 24OH and biomarkers of AD pathology and neurodegeneration in CSF samples from a memory clinic cohort, consisting of SCI, MCI and AD patients. More specifically, we were interested to explore the presence of sex-specific differences in these relationships. Demographic, clinical and biomarker data are shown in Supplementary Table 1. Women (n=42) and men (n= 48) did not differ in age. The diagnostic groups were similarly distributed between men and women, although SCI subjects were somehow overrepresented in women. CSF levels of Aβ42, t-tau, p-tau, NFL and 24OH were equally distributed across men and women. 24OH did not correlate with age (*p* = 0.951). Mean 24OH levels were not different between men and women (Figure Supplementary 2). To investigate associations between CSF 24OH and biomarkers of AD pathology and neurodegeneration, we performed separate linear regression models for Aβ42, t-tau, p-tau and NFL in all participants and in men and women, adjusting for age and diagnosis (Table 1). Increased 24OH levels were associated with decreased p-tau and NFL levels in the total cohort (β = −0.212, *p* = 0.041; β = −0.256, *p* = 0.001). When performing grouped analysis in men and women separately, the associations was seen in women (β = −0.294, *p* = 0.021; β = −0.230, *p* = 0.026) but not in men (β = −0.028, *p* = 0.865; β = −0.217, *p* = 0.082). In addition, higher 24HOC levels were associated with lower t-tau levels in women only (β = −0.235, *p* = 0.033). There was no correlation between 24OH and Aβ42 in the total population or in men and women analyzed separately.

**Table 1.** Associations between CSF 24OH and biomarkers of AD pathology and neurodegeneration. Associations of Aβ42, t-tau, p-tau and NFL with 24OH were examined using multiple linear regression models, adjusting for age and diagnosis. Data shown are β coefficients with *p* values (p<0.05 was considered significant and marked in bold). Models were tested in the whole cohort and in men and women separately.

| | **Aβ42**<br>$\beta$ (p-value) | **t-tau**<br>$\beta$ (p-value) | **p-tau**<br>$\beta$ (p-value) | **NFL**<br>$\beta$ (p-value) |
| --- | --- | --- | --- | --- |
| *All participants* |  |  |  |  |
| 24OH | .030 (.628) | -.085 (.324) | -.212 (**.041**) | -.256 (**.001**) |
| *Men* |  |  |  |  |
| 24OH | -.043 (.670) | .121 (.365) | -.028 (.865) | -.217 (.082) |
| *Women* |  |  |  |  |
| 24OH | .060 (.712) | -.235 (**.033**) | -.294 (**.021**) | -.230 (**.026**) |

## DISCUSSION

CYP46A1 regulation in the brain has received increasing attention as possible therapeutic target for neurodegenerative diseases. We have previously reported that CYP46A1 overexpression appears to be beneficial for aging in old female mice ^4^. These results were in agreement with previous studies showing positive effects of CYP46A1 up-regulation in female mouse models of AD ^9–11^. In the present study we aimed to deepen our understanding of CYP46A1 function in the brain by investigating the effects of its overexpression in male and female aged mice. Aging is known to affect behavioral performances in wild type mice by decreasing spontaneous alternations in Y- maze test, the exploration time in lit compartment of the dark light box paradigms and worsening spatial learning ^35^. We show that CYP46A1 overexpression in female mice positively modulates anxiety-like behavior in Dark light box test and long and short term hippocampal-dependent memory in MWM test^4^ and Y-maze tests, protecting against the normal deleterious effects of aging on these parameters. In contrast, males overexpressing CYP46A1 displayed a significant worsening of spatial-dependent memory retention and anxiety-like behavior when compared to their control littermates. These sex-specific differences were not seen in young Cyp46Tg, suggesting that the effects of CYP46A1 overexpression arise along with aging, conferring protection for females and further deterioration for males.

Given the association of behavioral paradigms with hippocampal function, we further investigated whether hippocampal dendritic spines from Cyp46Tg and WT mice displayed morphological differences, underlying a possible correlation with the observed phenotypes. During aging, the morphology of dendritic spines changes and spine density and total number of spines diminishes, both in cortical and hippocampal region including CA1^36, 37^. These changes have been observed in AD patients ^37, 38^ and in AD mouse models, where a lower spine density is preceded by reduction of spine length and neck enlargement^39^. Old Cyp46Tg males displayed lower spine density when compared to age-matched littermates. Furthermore, the decrease of spine density seen in old male Cyp46Tg mice can be related to increased anxiety-like behavior, as anxiety phenotype in wild-type mice has been previously characterized by decreased spine density in CA1 regions ^40^. In old Cyp46Tg females, while the spine density was not significantly altered, longer and larger spines were shown. Larger dendritic spine heads correlate with augmented AMPA receptors in post synaptic densities, and with neurotransmission improvements relevant for Long-Term Potentiation and memory processes ^41–43^. Thus, the morphological features of dendritic spines from old female Cyp46Tg support the enhancement of hippocampal memory and the higher levels of synaptic markers ^4^ observed in these mice.

Differences in 24OH levels cannot account for the sex-induced difference seen in CYP46A1 overexpressing mice as both males and females show similar amounts of 24OH in brain ^4^. 24OH has been reported as allosteric modulator of NMDAR ^19^ and enhancement of NMDAR signaling has been observed in the hippocampus of both Cyp46Tg males and females compared to control mice. Thus, it is unlikely that a differential upregulation of NMDAR signaling could be part of the mechanism by which CYP46A1 overexpression induces sex-differential effects on cognition during aging.

Cholesterol is the precursor of steroidal hormones in the periphery and also in the brain, where neurosteroidogenesis takes place^33^. Cholesterol metabolites have been investigated as possible ligands of estrogen receptors^23, 44^. We therefore explored whether the sex-specific differences observed in our study could be mediated by brain estrogen signaling. Cyp46Tg females showed increased *Cyp19a1* and *Hsd17b1* mRNA levels (correlating with higher activity of the enzyme ^45^), suggesting an increase in local production of E2 in the brain. The higher levels of aromatase in hippocampus from Cyp46Tg females did not correspond to higher levels of E2 compared to WT controls, although E2 hippocampal levels were more homogenous among Cyp46Tg females. E2 is highly lipophilic and can easily pass from the periphery to the brain. In female mice older than 13 months, the reduction of oocyte follicles and the consequent decrease in blood E2 levels ^46^ could affect brain E2 status, that would depend mainly on brain local production, compromising E2-dependent neuroprotective mechanism ^47^. The concept of an enhanced E2 signaling in the brain of Cyp46Tg females is also supported by the fact that *Ers2* and *Apod*, an ERβ-specific target gene ^48^, are increased in the hippocampus of these mice. Dendritic spines from memory-related regions as hippocampus are highly sensitive to estrogen signaling and spine density has been found to be proportional to estrogen levels ^49–52^. Moreover, exogenous estrogen increases the number of mushroom shaped spines and levels of postsynaptic markers in CA1 region in mice ^53^. We describe that 24OH itself increases ERβ estrogen receptor levels in hippocampal neurons and furthermore, activates both the estrogen non-genomic mediated rapid signaling (by inducing CREB phosphorylation) and the estrogen genomic pathway by inducing nuclear translocation of ERβ, where the transcription of ER gene targets as *Bdnf* is thus promoted. Noteworthy, 24OH also increases Arc levels in hippocampal neurons. Expression of Arc is required for BDNF induced LTP and is linked to estrogen activity ^54, 55^. A sustained activation of brain estrogen signaling by 24OH could in turn promote neuroprotection and enhance cognitive functions in in Cyp46Tg females during aging, acting as replacement for the loss of endogenous E2. Together our results suggest that the enhancement of estrogen receptor signaling in Cyp46Tg females could be a consequence of a 24OH dependent-neurosteroid activation in the brain. 24OH is a potent agonist for LXR and activation of LXR has been reported to increase expression of neuroactive steroids in male rats, for example increasing mRNA of P450scc and 5α-Reductase ^22^.In contrast to females, Cyp46Tg males did not show enhancement of estrogen receptor signaling in hippocampus. Nevertheless, enzymes as *Srd5a1, Hsd17b10* contributing to the formation of DHT, as well as DHT levels, were significantly increased. High concentrations of DHT could contribute to the reduction of CA1 spine density, worsening of memory performance and anxiety like behavior in Cyp46Tg males. According to previous finding, aromatase inhibition, and thus, the conversion of testosterone into E2, was shown to cause dendritic spine loss in mice ^56^. Further, testosterone and its metabolites were reported to have anxiolytic effects in male mice ^57, 58^. Finally, our in vitro results suggest that excessive DHT levels in Cyp46Tg males counteract the positive effects exerted by 24OH on estrogen signaling and behavior.

LXR are involved in multiple signaling pathways. When the dimer LXR-RXR is formed, it binds to the LXR-RE promoter, inducing the transcription of gene targets as *Srebf1* and promoting Retinaldehide dehydrogenase (Raldh) activity ^59, 60^. Raldh in turn synthesizes all-*trans*-retinoic acid which can activate RARs. We show that 24OH increases RARα levels and activity in hippocampal neurons, as seen by enhanced mRNA and protein levels of the nuclear receptor as well as *Cyp26b1*, the enzyme responsible for retinoic acid catabolism ^61^. We also show that DHT is able to revert these effects. RARα can be located at synapses and participates to higher cognitive functions in hippocampus and has a key role in spine formation and dendritic growth ^62 63^. Importantly, RAR signaling promotes the expression of target genes involved in neurosteroidogenesis in the periphery ^64^. Therefore, RAR activation could contribute to the mechanisms by which 24OH regulates neurosteroid metabolism. Moreover, gene expression of ADAM10 is dependent on RARα signaling ^34^. Among many other substrates, ADAM10 cleaves APP into soluble APPα associated resulting in a non-amyloidogenic pathway ^65^. APPα was recently reported to promote neuroprotection and enhance learning and memory functions and neuronal structural integrity ^66^. Noteworthy, levels of ADAM10 were found higher in hippocampus of female Cyp46Tg as compared to female littermates, while unchanged between male groups. Enhanced ADAM10 activity, leading to increased production of soluble APPα and consequent neuroprotection, could also participate in beneficial effects of 24OH in Cyp46Tg females.

Finally, when analyzing the associations between 24OH levels and markers of neurodegeneration in CSF from patients in a memory clinic cohort, we could see that higher 24OH was associated to lower tau, p-tau and NFL levels in women but not in men. This association was seen regardless of AD pathology and with comparable CSF levels of 24OH in men and women. Higher 24OH levels in men did not correlate with higher p-Tau and NFL. These findings suggest that the sex specific functions of 24OH observed in female mice in our study may be translated mainly to women and future studies in larger cohorts will be necessary to further assess how levels of 24OH affects AD pathology in both sexes. Importantly, in the present cohort levels of 24OH did not differ in AD cases when compared to control and MCI as previously reported^14–16^. In this context, it is important to mention that the patients selected in this cohort were not users of medications for hypercholesterolemia, hypertension or diabetes, known to affect cholesterol metabolism.

In summary, our findings in mice supports the hypothesis that CYP46A1 overexpression and higher levels of 24OH modulate neuroactive steroid signaling in the brain in a sex-dependent manner, leading to different outcomes on cognitive functions and neurodegenerative processes in males and females. This is particularly important from a clinical perspective, considering that a higher number of women suffer from AD and they are subjected to faster progression rates from MCI to AD than men. Moreover, early menopause before the age of 40–45, resulting from ovarian removal, premature ovarian insufficiency, chemotherapy and aromatase inhibitor treatment; represents a female-specific risk factor for higher cognitive decline and higher levels of AD neuropathology ^67–69^. Abrupt loss of ovarian hormones, including E2, is therefore associated to age-related pathophysiological processes ^70^. In light of these results, CYP46A1 activation could represent a pharmacological target for modulation of neurosteroidogenesis that could specifically enhance brain E2 signaling in women at risk of developing AD. Based on our data, we suggest that targeting CYP46A1 for therapy of AD or other neurodegenerative diseases may lead to different outcomes in men and women. Finally, this study highlights the importance of taking into account the sex dimension in both preclinical and clinical studies of neurodegenerative diseases like AD.

## ACKNOWLEDGEMENTS

This research was supported by Margaretha af Ugglas Stiftelse, National Institute On Aging of the National Institutes of Health under Award Number R01AG065209, the regional agreement on medical training and clinical research (ALF) between Stockholm County Council and Karolinska Institutet 592522, Swedish Research Council, Gun och Bertil Stohnes Stiftelse, Karolinska Institutet Foundation for geriatric research, Stiftelsen Gamla Tjänarinnor and Tore Nilsson Stiftelse. Work in Swansea was supported by the Biotechnology and Biological Sciences Research Council (BBSRC, grant numbers BB/S019588/1 to WJG, BB/L001942/1 to YW) and the European Union through European Structural Funds (ESF).

The behavioral studies in mice were performed at the Animal Behavior Core Facility (ABCF) of Karolinska Institutet.

## CONFLICTS OF INTEREST

WJG and YW are listed as inventors on the patent “Kit and method for quantitative detection of steroids” US9851368B2 licenced by Swansea University to Avanti Polar Lipids Inc and Cayman Chemical Company. WJG and YW are shareholders in CholesteniX Ltd.

## FIGURE LEGENDS

**Figure S1.**
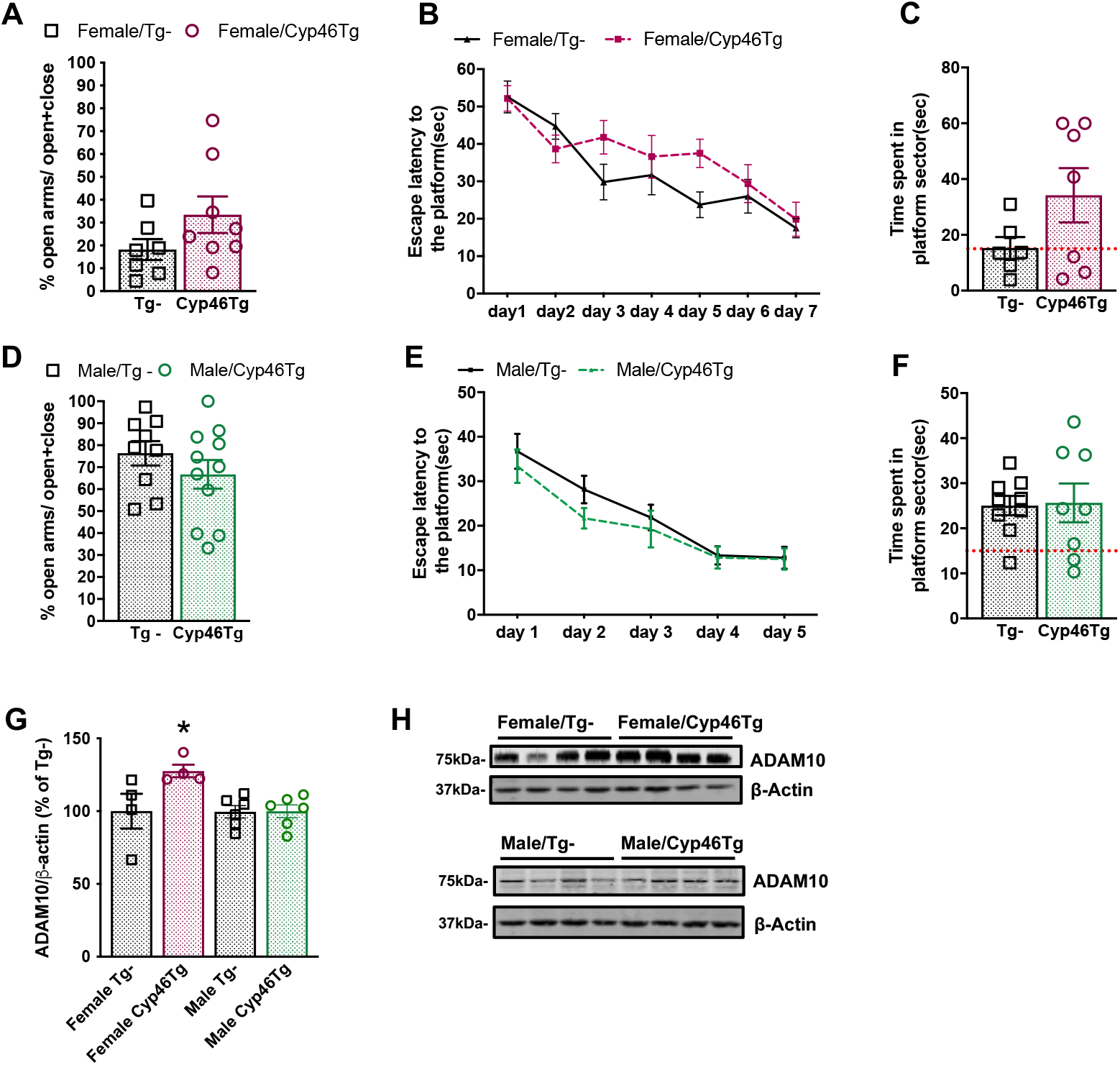
No differences in EPM and MWM tests were found in adult Cyp46Tg male and female mice. Behavioral test were performed in 9 months old Cyp46 Tg female (upper part, burgundy graphs) and male mice (lower side, green graphs) **(A-F,** N=7-11 mice/group/sex). Percentage of time spent in open arms over the 5 min duration of the EPM test **(A**, **D)**. Escape latency (time required to reach hidden platform) over 5 days acquisition phase in MWM test in Cyp46 Tg female and male mice **(B, E)**. Time spent in the quadrant where the platform was located during MWM probe test after 24 hours (**C, F)**. **ADAM10 levels are increased in hippocampus from Cyp46 Tg female mice, while unchanged in males.** Immunoblotting analysis of ADAM10 in 20 months old Cyp46 Tg male and female mice **(G, H**, *P<0.05, N=4-6 mice per group). Data are shown as mean ± SEM of immunoreactivity (OD × area of the band) normalized by actin levels.

**Figure S2.**
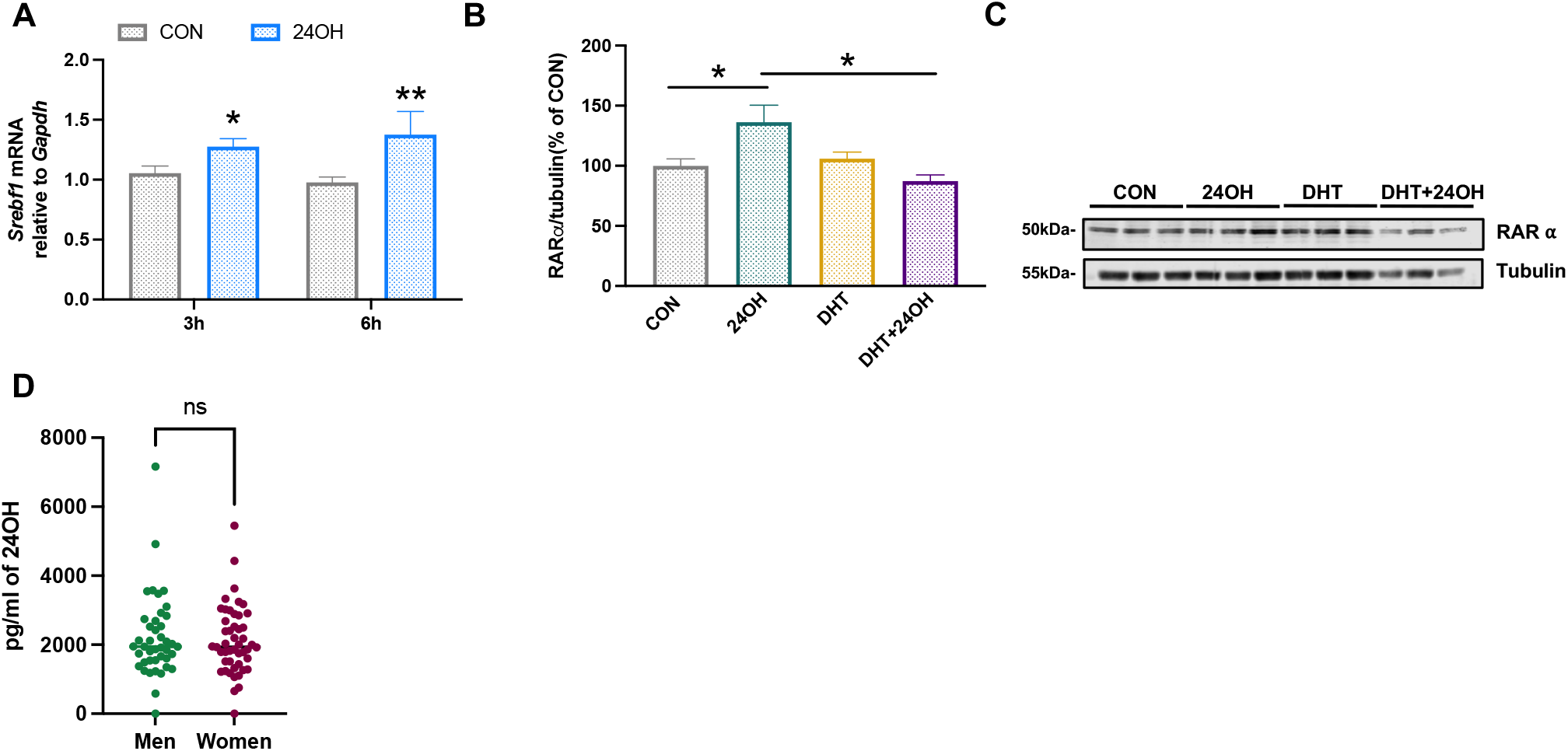
24OH *in vitro* treatment at different time points. *Srebf1* mRNA levels measured in primary hippocampal neurons after 6-hour and 12-hour treatment with either 1 μM of 24OH or its vehicle as a control **(A,** *P<0.05, **P<0.01, N=3 independent experiments, each in triplicate). Immunoblotting analysis **(B)** and representative blot **(C)** after 24 hour-treatment with 1 μM 24OH, 10 nM DHT and the combination of both in RARα levels in hippocampal neurons. Quantification was calculated by OD × area of the band and normalizes by tubulin levels (*P<0.01, N=3 independent experiments, performed in triplicated). **CSF 24OH levels are unchanged between men and women in a memory clinic cohort.** 24OH levels in CSF samples measured by Liquid Chromatography/Mass Spectrometry (LC/MS) **(D,** ns=no significant).

**Table Supplementary 1.** Demographic, clinical and biomarker data. Continuous data are shown as mean (± SD). Demographic factors, clinical characteristics, and CSF biomarkers were compared using χ^2^ and Mann-Whitney U tests.

| Variables | Men<br>(n= 42) | Women<br>(n= 48) | *p* value |
| --- | --- | --- | --- |
| *Demographics* |  |  |  |
| Age | 65.0 (± 6.5) | 65.8 (± 7.7) | .537 |
| *Diagnosis* |  |  |  |
| SCI | 12 | 18 | .669 |
| MCI | 15 | 15 |  |
| AD | 15 | 15 |  |
| *CSF biomarkers (pg/mL)* |  |  |  |
| Aβ42 | 683.3 (±228.4) | 722.8 (± 269.8) | .450 |
| t-tau | 397.2 (±196.5) | 444.1 (244.3) | .462 |
| p-tau | 67.7 (± 27) | 68.2 (± 22.7) | .442 |
| NFL | 1387.6 (±674.7) | 1330.8 (± 850.1) | .290 |
| 24OH | 2206.0 (±1194.9) | 2124.0 (± 980.0) | .913 |

## Notes

### Competing Interest Statement

William J. Griffiths and Yuqin Wang are listed as inventors on the patent: Kit and method for quantitative detection of steroids US9851368B2, licenced by Swansea University to Avanti Polar Lipids Inc and Cayman Chemical Company. WJG and YW are shareholders in CholesteniX Ltd.

